# ELAVL3 regulates splicing of RNAs encoding synaptic signaling proteins in D1 and D2 striatal medium spiny neurons

**DOI:** 10.1101/2024.11.18.622571

**Authors:** Krithi Irmady, Claudia Scheckel, Ruth A Singer, Thomas Carroll, Robert B Darnell

## Abstract

The neuronal RNA-binding protein (RBP) family nELAVL regulates key neuronal processes by binding directly to target RNA transcripts. In this study, we demonstrate that ELAVL3 is the predominant nELAVL paralog expressed in D1 and D2 medium spiny neurons of the striatum. To investigate its function, we developed ELAVL3 cTag-crosslinking and immunoprecipitation (CLIP) to generate RBP-RNA interaction maps from these neurons. By integrating data from ELAVL3-cTag and Elavl3 knockout mice, we identified distinct regulatory effects of ELAVL3 on alternative splicing of its target transcripts. Notably, ELAVL3 modulates splicing of transcripts encoding proteins critical for glutamate and dopamine receptor signaling. These findings underscore the role of ELAVL3 in RNA-mediated regulation of molecular pathways essential for medium spiny neuron function in the striatum.

## Introduction

RNA-binding proteins (RBPs) play crucial roles in regulating neuronal functions in the brain. RBPs control messenger RNA (mRNA) localization, levels, alternative polyadenylation, and splicing, therefore generating protein diversity from a limited genome^1^. This regulation is particularly important for key neuronal processes such as neurogenesis, synaptic activity, and plasticity^2^. Dysregulation of RBPs has been implicated in several neurodegenerative disorders, including frontotemporal dementia, Alzheimer’s disease, and amyotrophic lateral sclerosis^3–5^.

Neuronal ELAVL (nELAVL) proteins are a family of neuron-specific RBPs homologous to the *Drosophila* embryonic lethal abnormal vision (elav) gene. First identified in the context of paraneoplastic neurological diseases, these paralogs, ELAVL2, ELAVL3, and ELAVL4, are highly expressed in neurons^6^ and regulate RNAs via mechanisms including localization, stability and alternative splicing. We have previously demonstrated that neuronal ELAVL plays a pivotal role in synaptic signaling and postsynaptic processes in the mouse cerebral cortex^7^.

The striatum, a subcortical brain region, is crucial for cognitive and motor control. Medium spiny neurons (MSNs), which constitute over 95% of striatal neurons, integrate cortical and thalamic glutamatergic inputs with dopaminergic signals from the substantia nigra^8^. MSNs are divided into two subtypes, D1 and D2 neurons, which predominantly express D1 and D2 dopamine receptors respectively. Glutamate from corticostriatal and thalamostriatal projections activates MSNs, with D1 neurons (direct pathway) promoting movement and D2 neurons (indirect pathway) inhibiting movement. Dopamine further modulates these neurons by binding to D1 and D2 receptors and regulating the cyclic adenosine monophosphate (cAMP)-protein kinase A (PKA) pathway. D1 receptors, coupled to G-stimulatory proteins, enhance adenylyl cyclase and increase cAMP levels, while D2 receptors, coupled to G-inhibitory proteins, reduce adenylyl cyclase activity and cAMP levels^9,10^. Dopamine thus stimulates D1 neurons and suppresses D2 neurons, fine-tuning movement control. The regulation of cAMP in MSNs is further regulated by phosphodiesterase (PDE)s, which selectively degrade cAMP^11^. cAMP-PKA pathway in MSNs ultimately regulate maintenance of phosphorylation of proteins crucial for synaptic signaling^12,13^. Dysregulation of glutamate and dopamine receptor mediated pathways therefore disrupts synaptic signaling and impairs neuronal function in the striatum.

We have previously demonstrated that mRNA for the RBP ELAVL3 is highly expressed in the striatum, compared to other paralogs of the neuronal ELAVL family^7^. In this study, we investigated the role of ELAVL3 in RNA regulation within striatal MSNs. Using conditionally tagged ELAVL3 mice, we performed cross-linking and immunoprecipitation (CLIP) followed by sequencing to identify ELAVL3 RNA targets specifically in D1 and D2 MSNs. To assess the functional impact of ELAVL3 binding, we examined alternative splicing in the striatum from *Elavl3* knockout (KO) mice. Together, these findings demonstrate ELAVL3’s critical role in shaping RNA splicing patterns in MSNs and offer insights into the molecular mechanisms underlying synaptic signaling in the striatum.

## Results

### ELAVL3 is expressed in medium spiny neurons

We investigated expression of neuronal ELAVL paralogs by performing RNA-Seq in the mouse striatum. ELAVL3 was the highest expressed nELAVL paralog in the striatum, consistent with our previous work with in situ hybridization assays (**Fig. 1a**). We generated mice that express green fluorescent protein (AcGFP)-tagged ELAVL3 in a Cre-dependent manner (**Fig. 1b**). In order to maintain wild-type ELAVL3-RNA stoichiometry and regulation, we employed a knock-in strategy targeting the endogenous *Elavl3* locus (See methods). Conditionally tagged *Elavl3*-*AcGFP* (cTag) heterozygous and homozygous mice had no apparent phenotype. Immunofluorescence studies on *Elavl3* cTag mice bred with the ubiquitous cytomegalovirus (*CMV*)-Cre driver showed ELAVL3 protein expression in the striatum (**Fig. 1c**). To investigate the expression and function of ELAVL3 in striatal neurons, *Elavl3* cTag mice were bred with *DRD(D)1*-Cre and *DRD(D)*-2-Cre mice. Immunofluorescence confirmed ELAVL3 protein expression in D1 and D2 MSNs of the striatum and was consistent with known expression of *DRD1* and *DRD2* in the striatum in these mice^14,15^ (**Fig. 1d**).

**Figure 1.**
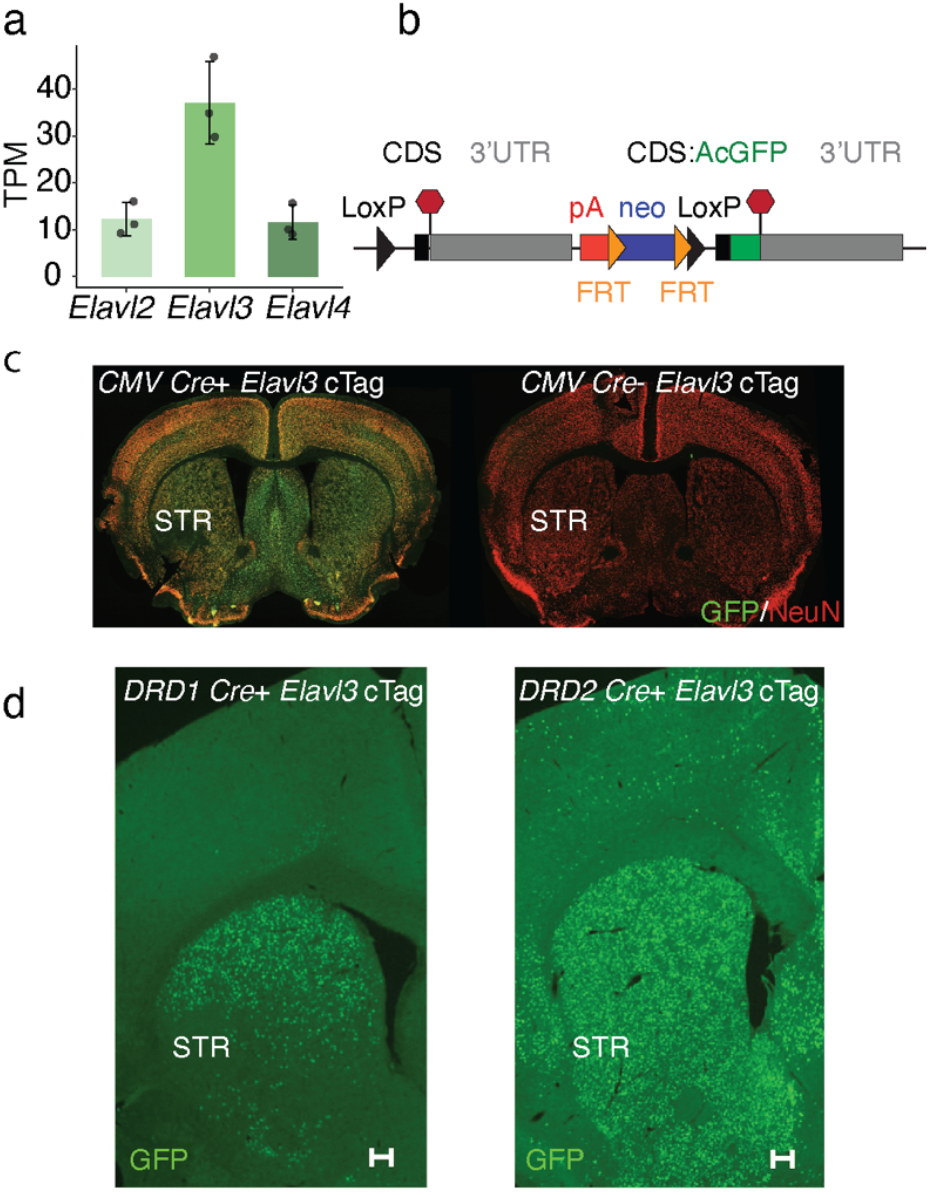
ELAVL3 is the predominant nELAVL paralog expressed in D1 and D2 MSNs of the striatum. (a) *Elavl3* is the highest expressed paralog of nELAVL family in the mouse striatum (n=3, wild-type mice) TPM: Transcripts per million (b) Schema of ELAVL3-AcGFP (*Elavl3* cTag) mouse illustrating a knock-in approach. The terminal exon and 3’UTR is flanked by loxP sites and AcGFP tagged terminal exon/3’UTR is inserted downstream of the transcription stop site (pA). CDS: Coding sequence, UTR: Untranslated region FRT: Flippase recognition target site (c) *Elavl3* cTag mice crossed with an ubiquitous promoter *CMV Cre* illustrate that ELAVL3 is expressed in the striatum (STR) of adult mouse brain. NeuN: Neuronal nuclei (d) *DRD(D)1* and *DRD(D)2* cre drivers crossed with *Elavl3* cTag mice illustrate that ELAVL3 is expressed in D1 and D2 MSNs in the striatum. Scale bar: 200 µm

### ELAVL3 directly binds to RNAs in D1 and D2 neurons

To identify the RNA targets of ELAVL3 in MSNs, we crosslinked cTag-ELAVL3 to its targets and performed GFP immunoprecipitation followed by sequencing (ELAVL3 cTag CLIP) in 2 month old mice expressing Cre drivers specific for D1 (*DRD(D)1*-Cre) or D2 (*DRD(D)*-2-Cre) neurons (**Fig. 2a**). ELAVL3 cTag CLIP identified 44822 significant binding sites (peaks) across 7810 genes in D1 and D2 expressing MSNs from three biological replicates per genotype. In both types of neurons, the majority of the binding sites were on intronic (∼75%), followed by 3’UTR regions of target transcripts (**Fig. 2b**). Motif analysis further confirmed that ELAVL3 binds specifically to U-rich motifs characteristic of neuronal ELAVL targets^7,16^, confirming the binding specificity of cTag-ELAVL3 (**Fig. 2c**). ELAVL3 binding peaks in D1 and D2 neurons included RNAs encoding proteins important for synaptic signaling such as glutamate receptors (*GRIA2, GRIN2B, GRM1, GRM7*), cAMP-PKA pathway regulators (*PDE4B, PDE4D, PDE10A*) and associated anchoring and signaling components (*PDE4DIP, PPP3CA)*, and proteins important for synaptic plasticity (*APP, RIMS1, DLG4, CACNA1C*). Gene ontology (GO) analysis on ELAVL3-bound RNAs identified terms related to glutamate receptor signaling, synaptic signaling, dendrite development, protein phosphorylation, RNA splicing and locomotion to be some of the top enriched terms (**Fig. 2d**).

**Figure 2.**
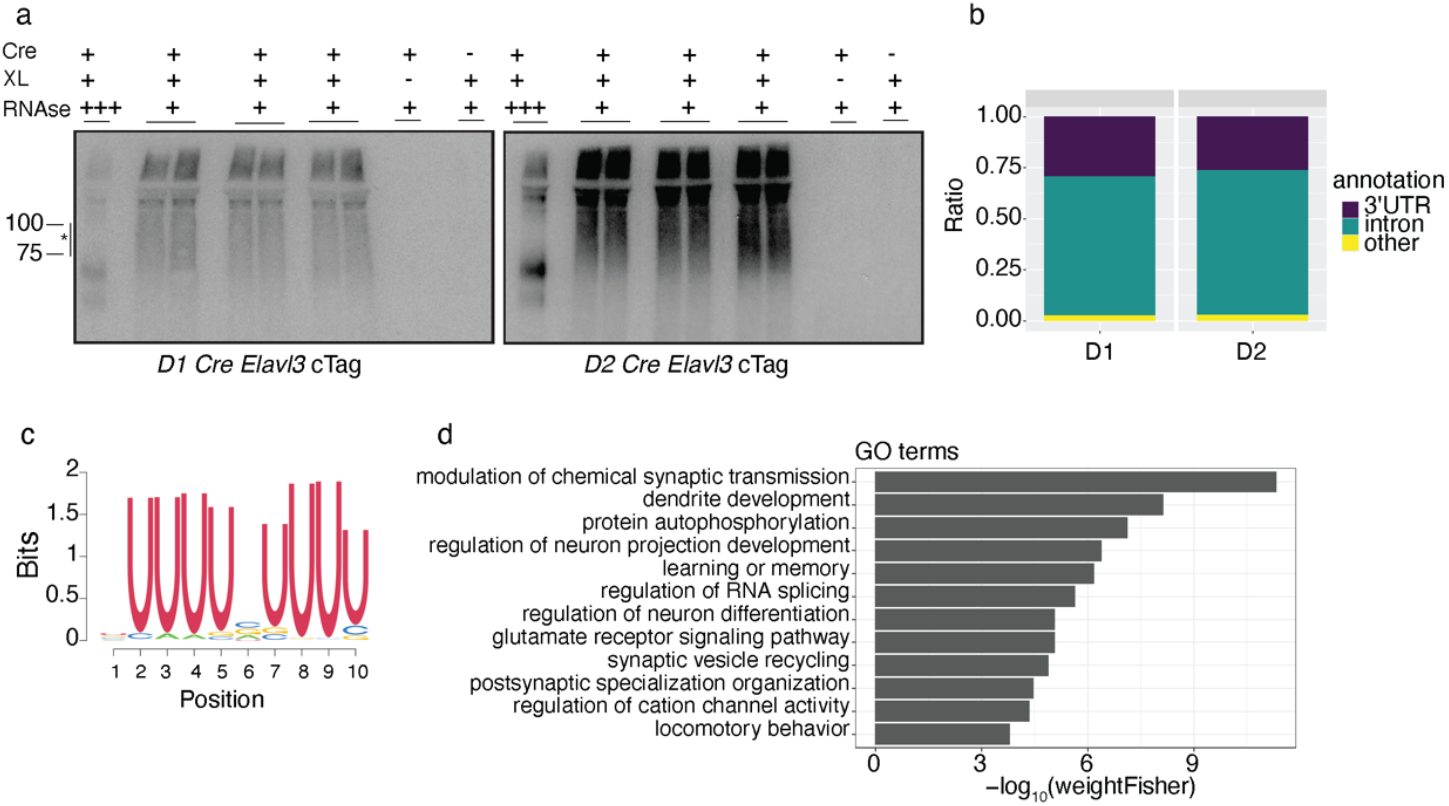
CLIP identifies ELAVL3 target transcripts in MSNs. (a) Representative autoradiograph for cTag CLIP shows ELAVL3-AcGFP (cTag):RNA complexes (*) pulled down upon crosslinking ELAVL3 to RNA in the presence of cre. No complexes are pulled down in the absence of *DRD(D1)1* or *DRD(D)2 cre* or in the absence of UV crosslinking. Striatum from 5-8 mice were pooled for each of the three replicates for *D1* and *D2* cTag CLIP. XL:UV crosslinking; +++:RNAse overdigestion; +:RNAse underdigestion (b) Genomic distribution of ELAVL3 binding on target transcripts show predominantly intronic and 3’UTR binding in D1 and D2 MSNs. (c) Motif analysis on significant ELAVL3 peaks identify U rich regions to be the preferred binding site for ELAVL3 (d) Gene ontology analysis on all significant peaks identify top GO terms associated with ELAVL3-bound transcripts.

We next assessed differences in ELAVL3 peaks between D1 and D2 neurons. Differential analysis identified 42 ELAVL3 peaks that were significantly different in D1 and D2 neurons (**Fig. 3a**). 24 peaks across RNAs from 10 genes were increased in D2 neurons compared to D1 neurons, while 18 peaks across 15 genes were increased in D1 neurons compared to D2 neurons. Differential peaks were noted in both the intronic (22 peaks) and 3’UTR (18 peaks) region. The most significant GO terms among the differentially bound RNAs were startle response, postsynaptic membrane assembly, glutamatergic synaptic transmission and calcium response (**Fig. 3b**).

**Figure 3.**
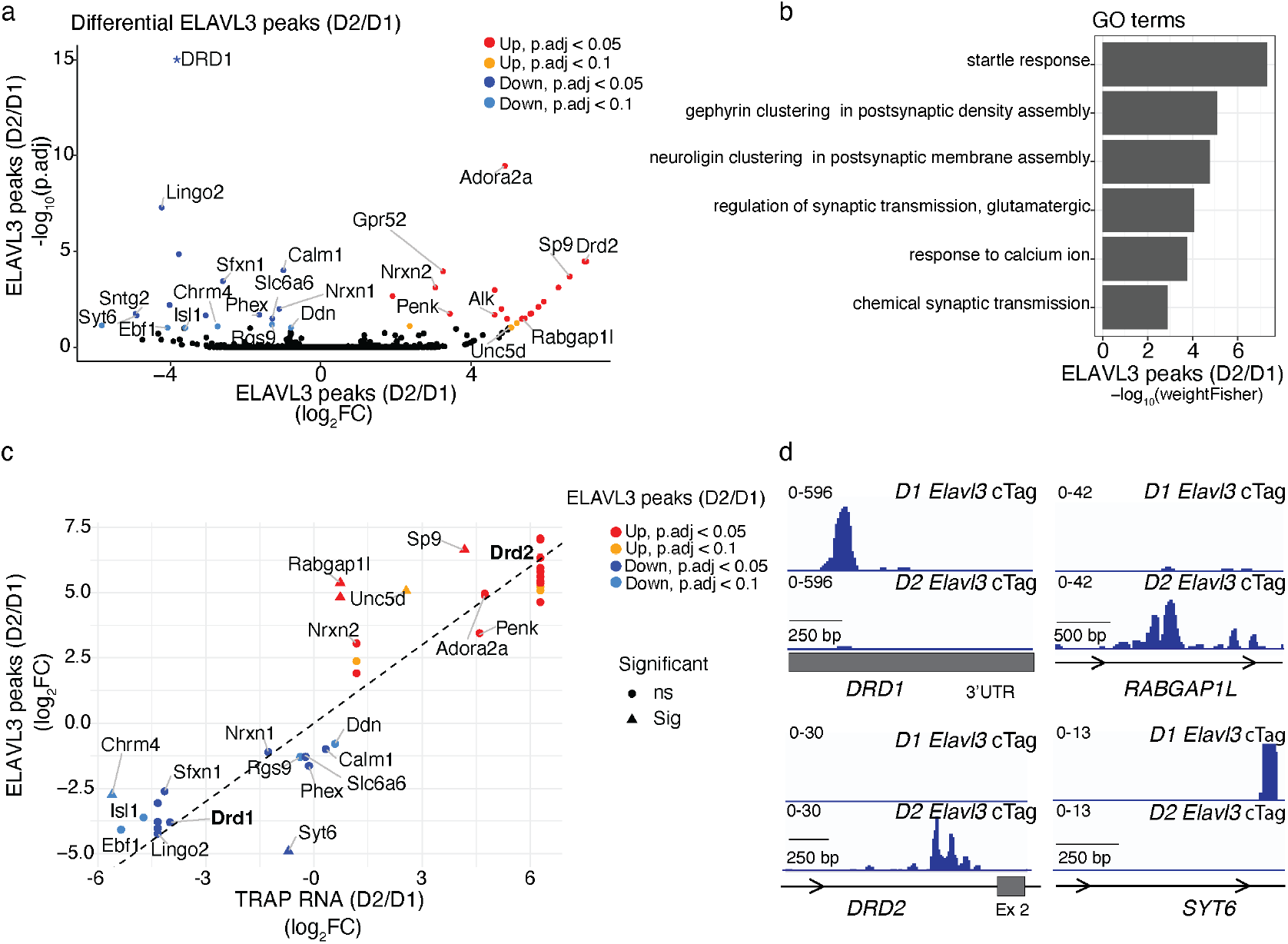
Differential binding of ELAVL3 in D1 and D2 MSNs. (a) Volcano plot shows differentially bound ELAVL3 RNAs identified by cTag CLIP in D1 and D2 neurons (n=3 mice per group). Each dot represents a peak (binding site). Differential peaks with p adjusted < 0.1 are highlighted according to direction of change. Only unique genes with peaks are labeled. * -log^10^ (p adjusted) > 20 (b) Top Gene Ontology terms associated with RNAs that are differentially bound in D1 and D2 neurons (c) Scatter plot illustrates fold change in ELAVL3 binding (CLIP, y axis) and RNA profile (TRAP, x axis) between D1 and D2 neurons. Peaks located greater than 1.96 standard deviations (95 percentile) from the slope are marked as significantly differentially bound (triangles), independent of differences in RNA profiles between cell-types. Only unique genes with peaks are labeled. (d) Genome viewer images show ELAVL3 CLIP peaks in D1 and D2 neurons on candidate transcripts that were differentially bound in D1 and D2 neurons.

Many of the differential peaks between D1 and D2 neurons corresponded to RNAs known to be differentially expressed between the two types of neurons. For example, in D2 neurons, ELAVL3 showed increased binding to *DRD2* RNA, consistent with its higher expression in D2 neurons (**Fig 3a**). Similarly, in D1 neurons, ELAVL3 binding was enriched on *DRD1* RNA, as expected given its specific expression in D1 neurons. These findings confirm that the observed differential binding aligns with the known cell-specific expression of these transcripts, validating the technical robustness of our approach and supporting the biological relevance of ELAVL3’s selective RNA interactions.

To further investigate the preferential binding of ELAVL3 in each MSN subtype, we examined its interaction with additional transcripts. Since ELAVL family members are known to regulate mRNA stability^17^, we first analyzed ELAVL3 binding independent of RNA abundance in each cell type. In D2 neurons, ELAVL3 showed increased binding to *ADORA2A, GPR52* and *PENK* RNAs, all of which are preferentially abundant in D2 neurons^18^ (**Fig 3a**). Similarly, in D2 neurons, ELAVL3 preferentially bound to *NRXN2, SP9, GPR52* and *RABGAP1L* RNAs. Conversely, in D1 neurons, we observed preferential ELAVL3 binding to *CALM1, CHRM4, SNTG2, NRXN1, RGS9, SFXN1, LINGO2* and *SYT6* RNAs. The proteins encoded by these transcripts are closely related to striatal neuronal functions or are of disease relevance (see Discussion).

To gain deeper insights into ELAVL3 binding specificity, we further examined mRNA profiles in the cell-types obtained by translating ribosome affinity purification (TRAP) approach in mouse D1 and D2 neurons^19^ as a proxy for mRNA abundance in each cell type. This confirmed that the differential binding of ELAVL3 to *DRD1* and *DRD2* RNAs reflected the differential abundance of these transcripts in D1 and D2 neurons respectively (**Fig. 3c,d**). For select RNAs (e.g., *RABGAP1L, SP9, UNC5D, SYT6, CHRM4*), ELAVL3 binding exhibited preferential binding in MSN subtypes that exceeded what would be expected based on differences in mRNA abundance between these neurons (**Fig. 3c,d**). Collectively, these findings identify RNAs that are both commonly and preferentially bound by ELAVL3 in D1 and D2 striatal MSNs, suggesting the potential for ELAVL3 to differentially regulate synaptic signaling in each cell type.

### Alternative splicing of RNAs in the striatum with loss of ELAVL3

Given that ELAVL3 bound predominantly to intronic regions in the striatum and are known to modulate RNA splicing of their target transcripts, we investigated RNA splicing differences in the striatum of 2 month old wild-type (n=3) and *Elavl3* knockout (n=3) striatum. In the *Elavl3* KO striatum, 1387 differential splice events were identified as significantly altered (**Fig. 4a**). These included skipped (cassette) exons and mutually exclusive exons (**Fig. 4b**). GO analysis of RNAs with significant splicing changes identified enrichment in terms related to glutamate receptor signaling pathway, G protein-coupled receptor signaling pathway, cAMP-mediated signaling, dendrite organization, and RNA splicing (**Fig. 4c**).

**Figure 4.**
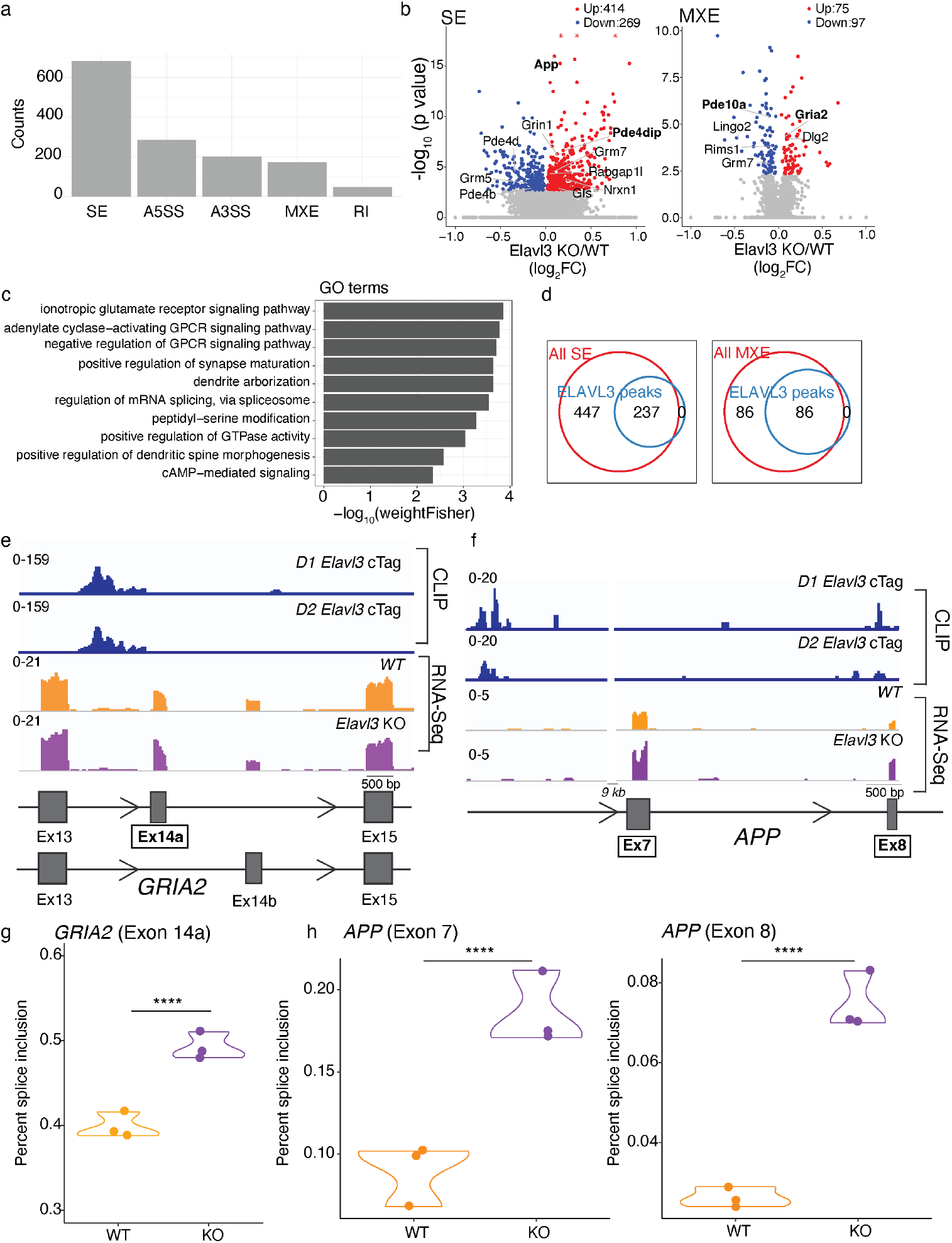
Alternative splicing in *Elavl3* knockout striatum. (a) Significantly spliced events in *Elavl3* knockout (KO) striatum (n=3) compared to wild-type mice (n=3). SE: Skipped (cassette) exon, A5SS: Alternative 5’ start site, A3SS: Alternative 3’ stop site, MXE: Mutually exclusive exon, RI: Retained intron. (b) Volcano plot shows significantly skipped and mutually exclusive exons in *Elavl3* KO (n=3) striatum compared to wild-type (n=3) mice. Each dot represents a splicing event. Significant events (FDR < 0.05) are colored according to direction of change. * -log10 (p value) >16 (c) Gene ontology analysis of RNAs that are significantly spliced in the *Elavl3* KO (n=3) compared to wild-type (n=3) striatum. (d) Venn diagrams illustrate the overlap between all significant splicing events (FDR < 0.05) in the *Elavl3* KO and splicing events with significant ELAVL3 peaks (FDR < 0.05) located in introns flanking the spliced exon. Only peaks detected in at least two out of three replicates in D1 or D2 neurons were included. e) Representative genome browser image illustrates ELAVL3 binding upstream of exon 14 in *GRIA2* RNA in D1 and D2 neurons. Increased inclusion of exon 14a is observed in the *Elavl3* KO mouse. (f) Genome browser image demonstrates ELAVL3 binding upstream of exons 7 and 8 in *APP* RNA. Increased inclusion of these exons is observed in the *Elavl3* knockout mouse. (g,h) Violin plots illustrate that exon 14 in *GRIA2*, and exons 7 and 8 in *APP* RNA shows increased inclusion in the *Elavl3* KO striatum compared to wild-type (WT) mouse (n=3 per group). **** p < 0.0001

To determine whether ELAVL3 directly regulates differential splicing of transcripts, we analyzed the presence of ELAVL3 CLIP peaks in intronic regions flanking skipped exons^7^. Of the 683 differentially skipped exons, 237 (∼35%) had a significant ELAVL3 intronic peak (detected in at least 2 of 3 replicates in either D1 or D2 neurons) located directly upstream or downstream, with some regions showing multiple ELAVL3 peaks (**Fig. 4d**). Similarly, in the *Elavl3* knockout striatum, 86 of 172 (∼50%) significant splicing events involving mutually exclusive exons were associated with an ELAVL3 binding site (detected in at least 2 of the 3 replicates in D1 or D2 neurons) in the flanking intronic regions of the spliced event (**Fig. 4d**).

Transcripts in which ELAVL3 differentially regulated splicing encoded proteins critical for glutamate receptor signaling (*GRIA2, GRM5, GRM7*) and phosphodiesterases, along with their interacting partners involved in cAMP-PKA pathway downstream of dopamine receptor activation (*PDE10A, PDE4B, PDE4D, PDE4DIP*). Other ELAVL3 target RNAs differentially spliced included those encoding proteins important for synaptic signaling and plasticity, including *APP, GLS, DLG1, DLG2, RIMS1*, and *ATXN1*. For example, in *GRIA2*, which encodes the GluA2 subunit of the AMPA receptor, ELAVL3 binding upstream of exon 14 was associated with increased inclusion of exon 14 in the *Elavl3* knockout compared to wild-type mice (**Fig. 4e,g**). This is consistent with previous findings that RBP binding upstream of an exon often correlates with decreased exon inclusion^7,20,21^. Other RNAs with differential splicing and ELAVL3 peaks in intronic region included *PDE10A*, a cAMP phosphodiesterase known to have the highest expression in MSNs^22^ and *PDE4DIP*, a phosphodiesterase 4D interacting protein (**Fig. S1**). RNAs differentially bound by ELAVL3 in D1 and D2 neurons, including *DRD1* and *DRD2*, did not show a difference in expression levels or splicing (**Fig. S2)**.

We and others have previously shown that binding of nELAVL paralogs to amyloid precursor protein (*APP*) RNA in the cerebral cortex promotes skipping of exons 7 and 8, generating *APP695*, the predominant neuronal isoform^16,23^. Consistent with this, we observed that ELAVL3 binds to *APP* RNA in D1 and D2 MSNs, and loss of ELAVL3 in the striatum is associated with increased inclusion of exon 7 and 8 in *APP* RNA (**Fig. 4f,h**). Together, these findings highlight the role of ELAVL3 in regulating splicing of its target transcripts, with implications for glutamate and dopamine receptor signaling and synaptic plasticity in the striatum.

## Discussion

In this study, we demonstrate ELAVL3 expression in D1 and D2 MSNs and investigate its RNA binding activity and functional effects in the striatum. We show that ELAVL3 binds to and impacts alternative splicing of several RNAs encoding proteins important for synaptic signaling and plasticity in the striatum.

Synaptic signaling in the striatum plays a central role in the physiological control of movement and cognition. Impaired activity of D1 and D2 neurons is a hallmark of striatal dysfunction in neurodegenerative diseases such as Parkinson’s disease and Huntington’s disease^24,25^. Furthermore, abnormal synaptic plasticity in the striatum contributes to treatment-related motor complications, including levodopa-induced dyskinesia in Parkinson’s disease patients^26^. ELAVL3 is an established regulator of synaptic signaling and synaptic plasticity in the cortex^7^. Our findings that ELAVL3 is expressed in D1 and D2 neurons of the striatum and binds RNAs encoding key proteins involved in synaptic signaling and plasticity suggest important roles for ELAVL3 in both normal striatal function and striatum-associated disease states.

Although ELAVL3 binds similarly to many transcripts present in both D1 and D2 neurons, its cell-type-specific roles may be influenced by the selective expression of its target transcripts within each neuron type, such as *DRD1* and *DRD2*. ELAVL3 targets the D2-enriched *PENK*, a proenkephalin--a precursor to enkephalins, opioid peptides implicated in regulating motor and emotional behaviors^27,28^. Notably, ELAVL3 binds to the GPCR *ADORA2A*, which activates the cAMP-PKA pathway upon binding to adenosine, and to *GPR52*, a GPCR known to modulate D2 neuron function, possibly through its interactions with *ADORA2A* and *DRD2*^29–31^. This suggests that ELAVL3 may modulate the GPCR activating pathways in D2 neurons.

In D1 neurons, preferential ELAVL3 binding was observed on *CHRM4*, a muscarinic acetylcholine receptor enriched in D1 neurons and known to influence cAMP signaling^32,33^. Additional targets preferentially bound by ELAVL3 in D1 neurons include RNAs with less defined roles in MSN subtypes, such as *CALM1*, which is involved in synaptic signaling and plasticity, and *LINGO2*, previously associated with Parkinson’s disease^34,35^. This suggests that ELAVL3 may modulate synaptic signaling in D1 neurons.

Other key ELAVL3 targets include the transcription factors *ISL1* and *SP9*, which are critical for differentiating MSNs into D1 and D2 subtypes, respectively^36,37^. ELAVL3 also binds to *NRXN1* and *NRXN2*, genes important for synapse assembly and pre- and postsynaptic signaling, with differential binding observed in D1 and D2 neurons^38^.

Interestingly, some targets demonstrated preferential ELAVL3 binding that exceeded RNA abundance differences. For instance, *SYT6*, a synaptotagmin family member involved in calcium-dependent vesicle fusion and neurotransmitter release, and *RABGAP1L*, which regulates GTPase activity and impacts mutant huntigtin protein levels^39,40^, exhibited ELAVL3 binding in D1 and D2 neurons respectively. *RABGAP1L* is notable for its epistatic effect on an another D2-enriched ELAVL3 target, *GPR52* which is encoded within an intron of *RABGAP1L*^40^.

A limitation of this study is that RNA profiles were not generated from the same cells used for CLIP, potentially complicating interpretations of ELAVL3’s effects. Although RNAs such as *DRD1* and *DRD2* did not show significant changes in expression or splicing in *Elavl3* knockout, other RBP functions, such as RNA localization and translation, which were not assessed in this study, may also be influenced by ELAVL3.

In this study, we found that ELAVL3 binds to numerous RNAs encoding proteins involved in postsynaptic signaling mediated by glutamate and dopamine receptors. In addition to the examples discussed above, ELAVL3 binds upstream of the differentially spliced exon 14 in *GRIA2*, a splicing event previously shown to be regulated by nELAVL paralogs in frontotemporal dementia^41^. This splicing event affects the dendritic localization of GluA2 and the signal transduction properties of AMPA receptors^42,43^. Similarly, ELAVL3 binds upstream of exon 6 in *PDE10A*, a striatal MSN-enriched phosphodiesterase, which is differentially spliced in the *Elavl3* knockout. *PDE10A* plays a critical role in MSN signaling by regulating the second messenger cAMP^44,45,11^. Notably, *PDE10A* expression is reduced in patients with Parkinson’s disease^22,46,47^, and drugs targeting this phosphodiesterase have also been explored in clinical trials for mood disorders^48^. Additionally, ELAVL3 binds to *PDE4B* and *PDE4DIP* RNAs, which also show differential splicing in the striatum of *Elavl3* KO mice. These proteins have been implicated in several neurodegenerative diseases, including Alzheimer’s disease, frontotemporal dementia and vascular dementia^49,50^.

We also observed that ELAVL3 binding on *APP* RNA was associated with increased skipping of exons 7 and 8 in the striatum in *Elavl3* KO mice, consistent with our previous work in the cortex^16^. *APP695*, the neuron-specific isoform, is characterized by the absence of exons 7 and 8, which results in the loss of the Kunitz-type protease inhibitor domain encoded by exon 7^51^. Beyond its role in amyloidogenesis, *APP* is critical for synaptic structure, function and plasticity, with *APP695* reported to influence glutamate receptor trafficking^52–54^. These findings reinforce the role of ELAVL3 in regulating alternative splicing events that affect proteins central to synaptic signaling and plasticity.

In summary, ELAVL3-driven differential splicing is expected to alter the expression and functionality of proteins that influence glutamate and dopamine receptor signaling, and synaptic plasticity with potential distinct effects in D1 and D2 neurons. Further studies are needed to determine the significance of specific ELAVL3 binding sites in the differential splicing of target exons and their contributions to RNA stability. Future investigations should also examine how ELAVL3-RNA interactions influence downstream signaling pathways that govern striatal function, movement, and cognition.

## Methods

### Mice

All animal procedures received approval from The Rockefeller University Institutional Animal Care and Use Committee (IACUC). *Elavl3* cTag mice were generated through standard restriction cloning similar to previous published studies for other RBPs^55,56^. *Elavl3* cTag targeting vector contains the endogenous *Elavl3* terminal exon preserving the 3’ UTR, triple synthetic poly(A) sites preceding *FRT-NEO-FRT* cassette, and terminal coding exon of *Elavl3* fused to AcGFP sequence. Construct was injected into B6 embryonic stem cells. The *FRT-NEO-FRT* cassette was removed by breeding heterozygous *Elavl3* cTag to FLPe expressing mice. Genotyping of *Elavl3* cTag mouse was performed at Transnetyx with standard assays for AcGFP and Neomycin. *CMV*-cre, *Drd1-cre* and *Drd2*-cre mice were obtained from commercial sources (Jackson laboratory, Mutant mouse resource and research center^14^*). Elavl3* KO mice were generated as described previously^7^.

### Immunofluorescence

Mice were perfused transcardially with phosphate buffered saline (PBS) followed by 4% paraformaldehyde (PFA) in PBS. Brains were dissected, fixed in 4% PFA in PBS at 4°C for 24 hours, immersed in 30% sucrose for 24 hours, and frozen before sectioning. Frozen brains were sliced into 30 μm thick sections on a cryostat (Leica). Brain sections were then subjected to immunofluorescence. Sections were washed, permeabilized, blocked with donkey serum and probed with primary antibodies for GFP (Invitrogen) and NeuN (Millipore) followed by incubation with Alexa 488 or 555 conjugated donkey secondary antibodies (Thermo Fisher Scientific). Sections were rinsed with PBS, mounted onto glass slides and cover slipped with Prolong Gold Antifade (Thermo Fisher Scientific). Imaging was performed on BZX700 (Keyence) microscope.

### cTag CLIP

cTag CLIP in 2 month old mice were performed using previously published protocols^55,56^. 5-8 striatum were pooled per genotype and experiments were performed in triplicate. For experimental controls, mouse brains that did not undergo UV irradiation, samples that underwent overdigestion, as well as brains from *Elavl3* cTag *Cre*(−) mice were used. Briefly, UV-crosslinked ELAVL3-AcGFP:RNA complexes were extracted using ice cold lysis buffer as described before^55^ and immunoprecipitated for 3 hours at 4 degree celsius using mouse monoclonal anti-GFP clones 19F7 (12.5 μg) and 19C8 (12.5 μg) mixture^55^ with Dynabeads™ Protein G magnetic beads (Thermo Fisher Scientific). Beads were then washed with a series of wash buffers as previously described^55^. RNA tags were dephosphorylated and ligated to an indexed degenerate 5’ RNA linker. RNA/protein complexes were eluted from beads and subjected to electrophoresis. Regions corresponding to 76-100 kDa were excised from the membrane and RNA tags were collected upon proteinase K digestion. The two-step polymerase chain reaction (PCR) amplification was performed utilizing reverse transcription primers with four-nucleotide barcode sequences for indexing. Samples were pooled together for HiSeq sequencing run (Illumina) and subsequently demultiplexed.

### RNA sequencing

RNA from mouse brains was Trizol (Invitrogen)-extracted, Ribo-Zero-selected (Epicentre, Madison, WI), DNase-treated (Roche), and prepared for sequencing, following the Illumina High-throughput TruSeq RNA Sample Preparation Guidelines. The libraries were sequenced for paired end reads on an Illumina HiSeq platform.

### Bioinformatics

CLIP reads were processed as described previously using CLIP Tool Kit (CTK)^57,58^. Briefly, raw reads were filtered for quality and demultiplexed using indexes introduced during the reverse transcription reaction. PCR duplicates were collapsed and adapter sequences removed. Reads were mapped to the mm10 RefSeq genome using BWA^59^. Mapped reads were further collapsed using fastq2collapse.pl from the CTK toolkit with default options. The distribution of CLIP reads that uniquely map to the genome was determined using bed2annotation.pl from the CTK toolkit. Uniquely mapped CLIP tags from all samples were pooled and CLIP peaks were called using tag2peak.pl from the CTK toolkit with a significance value cutoff of 0.05. For each sample, tags in significant peaks were counted using the “countOverlaps” R function. Peaks differentially bound by Elavl3cTag were determined by DESeq with a significance cutoff of p.adjusted < 0.1^58^. Differential gene expression for RNA-seq was performed using DESeq^60^. Splicing analysis was performed using RMATS with significant events called when FDR < 0.05^61^. Motif analysis was performed with MemeChip^61,62^. Gene ontology analysis was performed using TopGO (Alexa A, Rahnenfuhrer J (2024). *topGO: Enrichment Analysis for Gene Ontology*. R package version 2.58.0.).

## Acknowledgments

We extend our gratitude to Dr. Douglas Barrows and members of the Bioinformatics Resource Center, Genomic Resource Center, and Transgenic and Reproductive Technology core at Rockefeller University for their help and support. We thank Dr. Caryn Hale for help with bioinformatics and Ms. Paula Cutrim for technical assistance. We thank Dr. Nathaniel Heintz for sharing TRAP data^19^. This work was partially funded by the William N. and Bernice E. Bumpus foundation in support of KI. RBD is an Investigator of the Howard Hughes Medical Institute.

## Author contributions

Conceptualization, K.I., C.S., and R.B.D.; Data Acquisition and Analysis, K.I., C.S., R.A.S., and T.C.; Writing – Original Draft Preparation, K.I.; Writing – Review & Editing, K.I. and R.B.D. Funding Acquisition, K.I. and R.B.D.

