## Supplementary material for "ELAVL3 regulates splicing of RNAs encoding synaptic signaling proteins in D1 and D2 striatal medium spiny neurons": Fig S1, Fig S2

### Supplementary Figures

Fig S1

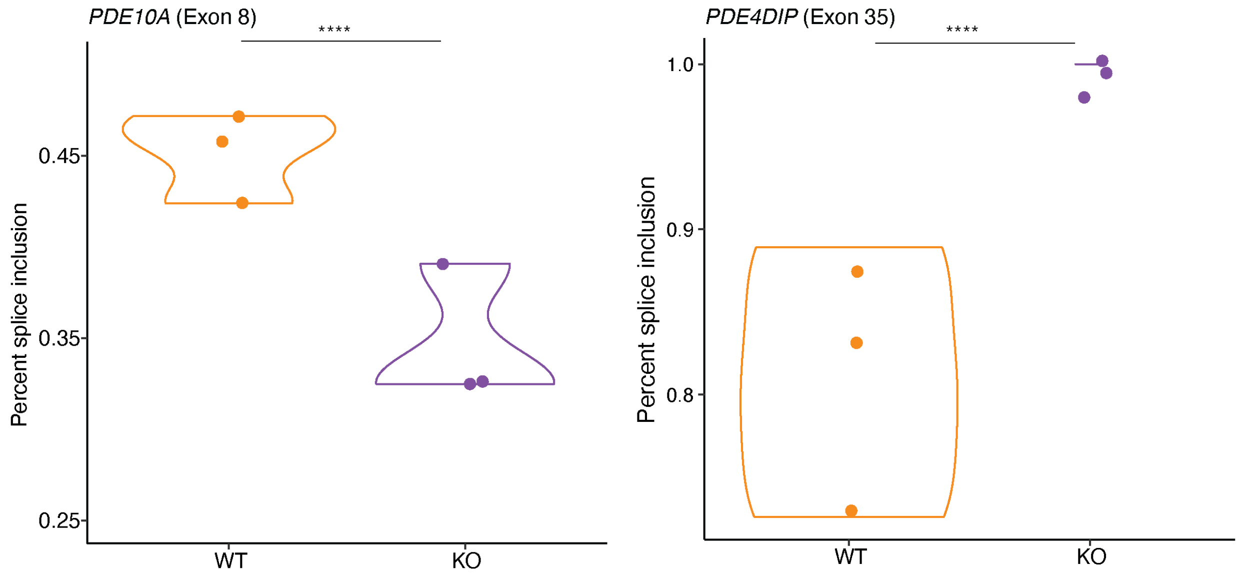


**Figure S1**. Significant splicing in *PDE10A* and *PDE4DIP* RNAs is observed in *Elavl3* knockout (KO) striatum compared to wild-type. n=3 per group. **** $p < 0.0001$

Fig S2

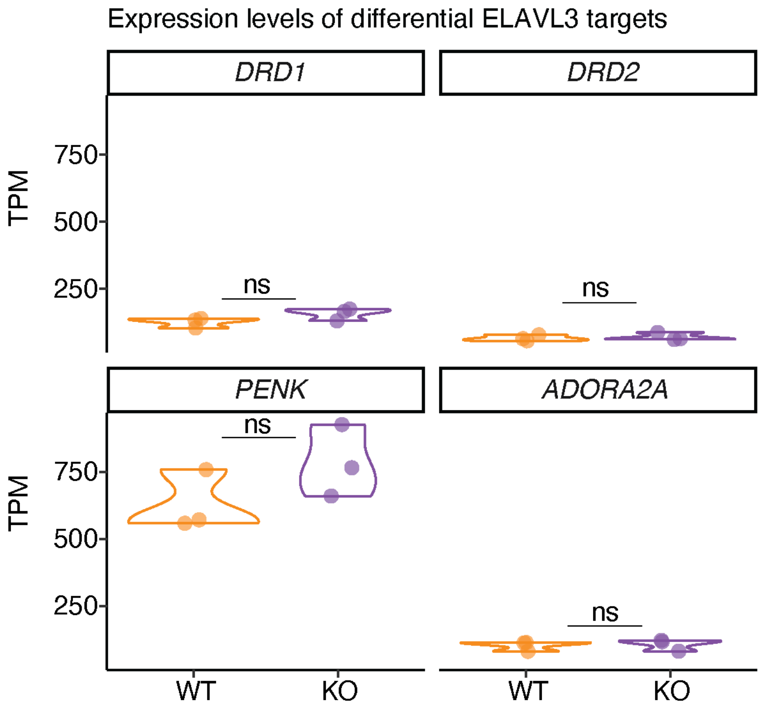


**Figure S2**. Expression levels of select RNAs differentially bound between D1 and D2 neurons in the striatum were not significantly changed in the *Elavl3* knockout (KO) compared to wild-type. n=3 per group. ns: not significant. p adjust values were obtained from DESeq analysis of differential gene expression between WT and *Elavl3* knockout striatum.
